# MTCH2 cooperates with MFN2 and lysophosphatidic acid synthesis to sustain mitochondrial fusion

**DOI:** 10.1101/2022.10.04.510812

**Authors:** Andres Goldman, Michael Mullokandov, Yehudit Zaltsman, Limor Regev, Smadar Zaidman, Atan Gross

**Affiliations:** Department of Immunology and Regenerative Biology, Weizmann Institute of Science, Israel

## Abstract

Mitochondrial dynamics is critical to sustain normal mitochondrial function and is linked to the response of cells to stressful conditions. Fusion of the outer mitochondrial membrane (OMM) is regulated by mitofusin 1 (MFN1) and 2 (MFN2), yet the differential contribution of each of these proteins to this process is less understood. Mitochondrial carrier homolog 2 (MTCH2) was shown to compensate for MFN2’s loss, however its exact function in mitochondrial fusion remains poorly understood. Here we determined the mitochondrial fusion-interplay between MFN1, MFN2 and MTCH2 and demonstrate that MFN2 and MTCH2 play separate, but redundant, roles required for mitochondrial fusion. Loss of either MFN2 or MTCH2 elicits mitochondrial fragmentation that retains mitochondrial plasticity, while loss of both proteins completely impairs mitochondrial fusion. We also show that expression of an MFN2 mutant targeted to the endoplasmic reticulum (ER) is sufficient to restore mitochondrial elongation in MTCH2 KO cells and that this restoration depends on the synthesis of the pro-mitochondrial fusion lipid lysophosphatidic acid (LPA). Moreover, silencing of MFN2 or inhibition of de novo LPA synthesis, revealed the requirement of MTCH2 to sustain mitochondrial plasticity in response to stress. Thus, we unmask two cooperative mechanisms that sustain mitochondrial fusion: one in the OMM, dependent on MTCH2 and MFN1, and independent of MFN2; and a second mechanism in the ER that relies on MFN2 and LPA synthesis.

## Introduction

Mitochondria are highly dynamic organelles that undergo continuous remodeling through fusion and fission events (Bereiter-Hahn and Vöth, 1994; Nunnari *et al*., 1997; Sesaki and Jensen, 1999; Rowland and Voeltz, 2012). Defects in mitochondrial fusion lead to fragmentation of the network into round shaped distinct mitochondria, while enforcing mitochondrial fusion leads to a highly interconnected network composed of elongated and tubular shaped mitochondria (Das and Chakrabarti, 2020)

Mitochondrial fusion relies on the activity of three large GTPases: optic atrophy protein 1 (OPA1), located in the inner mitochondrial membrane (IMM) (Olichon *et al*., 2002, 2003; Cipolat *et al*., 2004), and MFN1 and MFN2, located in the OMM (Santel and Fuller, 2001; Chen *et al*., 2003). MFN1 possesses higher GTPase activity compared to MFN2, and is essential for mitochondrial tethering during fusion (Ishihara *et al*., 2004). Although *in vitro* mitochondrial fusion assays showed that the formation of MFN1/MFN2 heterodimers yields the most efficient mitochondrial fusion rate (Meeusen, et al., 2004), MFN2 seems to be dispensable for mitochondrial fusion under different stimuli, such as amino-acid deprivation, stress-induced mitochondrial hyperfusion (SIMH) and inhibition of fission (Cipolat *et al*., 2004; Tondera *et al*., 2009; Rambold *et al*., 2011; Zhao *et al*., 2011; Lee *et al*., 2014). Besides its known role in mitochondrial fusion, MFN2 also localizes to the ER where it physically tethers mitochondria by interacting with MFN1/2 in the OMM (de Brito and Scorrano, 2008).

Mitochondrial carrier homologue 2 (MTCH2/MIMP/SLC25A50) is a non-classical OMM member of the mitochondrial carrier protein family (Robinson *et al*., 2012). Initially, MTCH2 was described as a mediator of apoptosis (Grinberg *et al*., 2005; Zaltsman *et al*., 2010) and later described to also regulate mitochondrial metabolism (Maryanovich *et al*., 2015; Buzaglo-Azriel *et al*., 2016; Ruggiero *et al*., 2017). More recently, it was described that MTCH2 is also involved in regulating mitochondrial dynamics (Bahat *et al*., 2018; Labbé *et al*., 2021). Several reports, which include genome wide association studies, and genetic models in *C. elegans*, zebrafish and mice, also demonstrated that MTCH2 plays a role in lipid metabolism (Kulyté *et al*., 2011; Bernhard *et al*., 2013; Bar-Lev *et al*., 2016; Landgraf *et al*., 2016; Rottiers *et al*., 2017; Labbé *et al*., 2021). MTCH2 deletion is associated with decreased lipid synthesis and storage (Buzaglo-Azriel *et al*., 2016; Landgraf *et al*., 2016; Rottiers *et al*., 2017), whereas increased MTCH2 expression was connected with increased lipid storage (Kulyté *et al*., 2011; Bar-Lev *et al*., 2016). In a recent study, MTCH2’s role in mitochondrial fusion was linked to the pro-mitochondrial fusion lipid lysophosphatidic acid (LPA), and it was proposed to funnel LPA towards the mitochondrial fusion machinery (Labbé *et al*., 2021).

LPA synthesis represents the first step of de novo phospholipid biogenesis (KENNEDY, 1961) and is formed by the addition of acyl-CoA to a glycerol-3 phosphate group, a reaction catalyzed by glycerol-phosphate acyl transferases (GPATs) (Coleman and Lee, 2004; Yamashita *et al*., 2014). Four different GPATs were described to date: GPATs 1 and 2, localized to the OMM (Coleman and Lee, 2004; Lewin *et al*., 2004; Pellon-Maison *et al*., 2007), and GPATs 3 and 4, localized to the ER compartment (Coleman and Lee, 2004; Jingsong *et al*., 2006). LPA synthesis is thought to primarily occur at mitochondria associated membranes (MAMS) (Pellon-Maison *et al*., 2007). Deletion of mitochondrial GPATs 1 and 2 elicits mitochondrial fragmentation, which can be rescued upon MFN2 overexpression (Ohba *et al*., 2013). In addition, reducing LPA synthesis using the GPAT inhibitor FSG67, which inhibits both ER and mitochondrial GPAT activity (Kuhajda *et al*., 2011; Clemens *et al*., 2019), also elicits mitochondrial fragmentation (Labbé *et al*., 2021).

Previously, we reported that MTCH2 deletion elicits mitochondrial fragmentation, which can be rescued by overexpressing MFN2, and that MTCH2 overexpression can restore mitochondrial elongation in MFN2 KO cells (Bahat *et al*., 2018). Here we report the occurrence of an unexpected functional complementation between MTCH2, MFN2 and LPA synthesis, which is necessary to sustain mitochondrial plasticity. MFN2 and MTCH2 are part of different but redundant/complementary pathways required to sustain normal mitochondrial architecture and plasticity. Loss of either MTCH2 or MFN2 does not impair mitochondrial fusion, whereas loss of both proteins abolishes the ability of mitochondria to fuse. We also show that targeting MFN2 to the ER is sufficient to restore mitochondrial elongation in MTCH2 KO cells, whereas inhibiting LPA synthesis suppresses MFN2’s rescue ability. Altogether, our results are consistent with a model where MTCH2, MFN2 and LPA synthesis cooperate and are necessary to sustain mitochondrial fusion in both steady state conditions and during cellular stress.

## Results

### MFN2 and MTCH2 compensate for each other’s absence

Previously, we showed that MTCH2 overexpression can restore mitochondria elongation in MFN2 KO cells, suggesting that MTCH2 can enforce mitochondrial fusion/elongation independent of MFN2 (Bahat *et al*., 2018). It was also recently shown that MTCH2 overexpression induces mitochondrial hyperfusion in HCT116 cells (Labbé *et al*., 2021). Since MTCH2 is not a dynamin-related protein (DRP), and thus lacks GTPase activity, its ability to induce mitochondrial fusion most likely depends on the presence of at least one of the MFNs.

To investigate this notion, we overexpressed MTCH2-GFP in WT, MTCH2 KO, MFN1 KO, MFN2 KO and MFN1/MFN2 double KO (DKO) MEFs. Overexpression of MTCH2-GFP elicits mitochondrial elongation in WT and MTCH2 KO MEFs and in approximately 20% of the cells it causes mitochondrial hyperfusion (Fig. 1A). Consistent with our previous report (Bahat *et al*., 2018), MTCH2-GFP overexpression enforces mitochondrial elongation in MFN2 KO but not in MFN1 KO and in MFN1/2 DKO MEFs (Fig. 1A, B). Thus, MTCH2-enforced mitochondrial elongation depends on the presence of MFN1 but not on the presence of MFN2.

**Figure 1.**
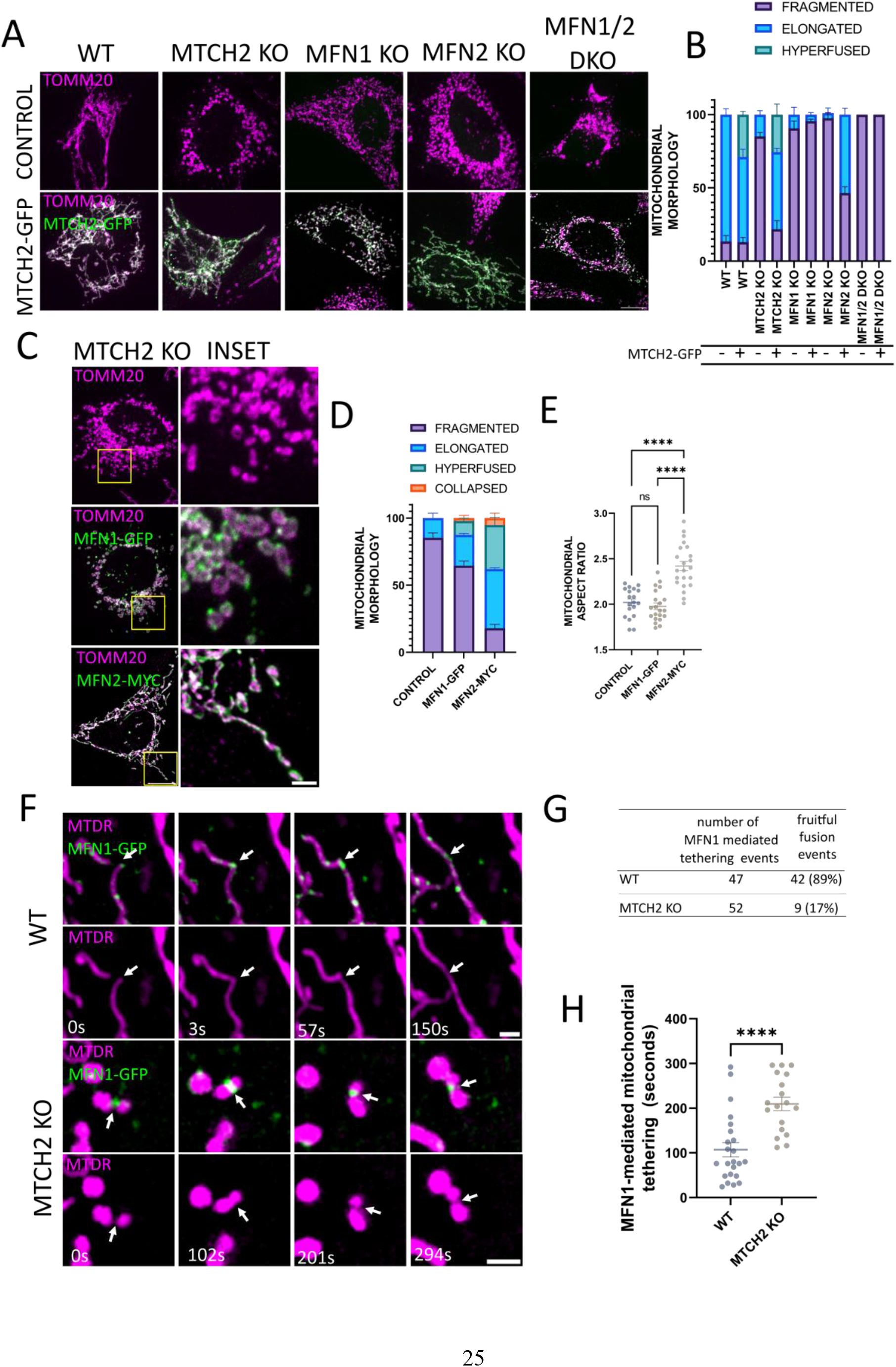
MFN2 and MTCH2 compensate for each other’s absence. **A** Representative images of WT, MTCH2 KO, MFN1 KO, MFN2 KO and MFN1/2 DKO MEFs either not expressing (control) or expressing MTCH2-GFP. Mitochondria was stained with αTOMM20 antibody. Scale bar 10 μm. **B** Classical classification of mitochondrial network morphology, showing average and SEM of three separate experiments (at least 20 cells were analyzed in per condition). **C** Representative images of MTCH2 KO MEFs either not expressing (control) or expressing MFN1-GFP or MFN2-MYC. Mitochondria were stained with αTOMM20 antibody. Scale bar 10 μm; inset 2 μm. **D** Classical classification of mitochondrial network morphology, showing average and SEM of three separate experiments (at least 20 cells were analyzed in per condition). **E** Mitochondrial aspect ratio analysis of MTCH2 KO either not expressing cells (control) or cells transfected with MFN1-GFP or MFN2-MYC (at least 20 control and transfected cells were analyzed per condition). One-way ANOVA statistical analysis. **F** Representative time lapse sequences of MFN1 mediated tethering events between mitochondria in WT or MTCH2 KO MEFs. Cells were transfected with MFN1-GFP and mitochondria were stained with mitotracker deep red (MTDR). Time lapse experiments were acquired for 5 minutes, capturing both channels every 3 seconds. Arrowheads follow a fruitful fusion event in WT cells, and in MTCH2 KO cells arrow heads depict an unfruitful fusion event, associated with long tethering time. Scale bar 2 μm. **G** Number of MFN1-GFP mediated tethering events monitored, and number of fruitful fusion events detected in WT and MTCH2 KO MEFs. **H** Quantification of MFN1-mediated mitochondrial tethering time in WT and MTCH2 KO MEFs. At least 20 MFN1-GFP-mediated tethering events were followed in WT and MTCH2 KO MEFs, according to the conditions described in F. The duration of the MFN1-GFP-mediated tethering event was determined by the time between the initial tethering and the lapse until mitochondria separated or fused with each other. Only tethering events that lasted more than 30 seconds were considered for this analysis to separate them from mitochondria “kiss and run” events. The data is presented as average tethering time and SEM, and statistical analysis was performed by t-test.

Previously, we showed that overexpression of MFN2 can induce mitochondria tubulation in MTCH2 KO cells (Bahat *et al*., 2018). Importantly, MFN1 was reported to have higher GTPase activity then MFN2 (Ishihara, Eura and Mihara, 2004), and thus we tested whether overexpression of MFN1 could also enforce mitochondrial fusion in MTCH2 KO cells. We generated an MFN1-GFP construct and validated its functionality together with MFN2-MYC by overexpressing them in MFN1 KO and MFN2 KO MEFs. As expected, MFN1-GFP rescued mitochondrial elongation in MNF1 KO MEFs, and MFN2-MYC rescued mitochondrial elongation in MFN2 KO MEFs (Supp Fig. 1A, B). As we previously reported (Bahat *et al*., 2018), MFN2 overexpression rescued mitochondrial elongation in MTCH2 KO MEFs, whereas overexpression of MFN1-GFP did not (Fig. 1C-E).

Next, we assessed whether MTCH2 deletion affected the expression levels of the mitochondrial fusion machinery proteins. Using both SDS-PAGE (Supp Fig. 1C) and blue-native gel electrophoresis (BNGE), we found no significant differences between WT and MTCH2 KO MEFs (Supp Fig. 1C, D). Moreover, OPA1 levels and isoform composition were unaffected by MTCH2 deletion (Supp Fig. 1C), suggesting that OPA1 cleavage or degradation is not driving MTCH2 KO dependent mitochondrial fragmentation.

To gain further understanding of mitochondrial fusion dynamics in MTCH2 KO cells, we performed live cell imaging studies of WT and MTCH2 KO MEFs expressing MFN1-GFP, and searched for mitochondrial fusion events marked by MFN1. In both WT and MTCH2 KO cells, MFN1-GFP was correctly targeted to the mitochondria and formed discrete foci (Fig. 1C and F). Importantly, we found successful mitochondrial fusion events associated with MFN1-GFP foci in WT cells (Fig. 1F, top timeline), which were rare in MTCH2 KO cells (Fig. 1G). In the absence of MTCH2, MFN1-GFP foci marked tethering events between mitochondria, but in most of the cases with no apparent resolution of the fusion intermediates (Fig. 1F, bottom timeline). To analyze MFN1-GFP associated tethering time, we analyzed tethering events between mitochondria that lasted more than 15 seconds, excluding “kiss and run” events from our analysis. MTCH2 deletion elicited close to a two-fold increase in MFN1-GFP associated tethering time between mitochondria (Fig. 1H), suggesting a defect in mitochondrial fusion probably downstream of MFN1 tethering.

### MFN2-targeted to the ER is sufficient to restore mitochondrial elongation in MTCH2 KO cells, and loss of MTCH2 results in increased ER-mitochondria contact sites

Next, we wanted to further understand how MFN2 compensates for the absence of MTCH2. Due to its dual function as both a mitochondrial fusion protein and an ER-mitochondria tethering factor, we investigated if MFN2’s ability to restore mitochondrial elongation in MTCH2 KO cells stems from its function at mitochondria or/and at the ER. To address this issue we expressed an MFN2-mito-targeted protein (MFN2-ACTA) (Annis *et al*., 2001) or an MFN2-ER-targted protein (MFN2-IYFFT) (Rojo *et al*., 2002) in MTCH2 KO cells.

In MFN2 KO MEFs expression of either MFN2-ACTA or MFN2-IYFFT resulted in a relatively small increase in mitochondrial tubulation, while co-expression of both proteins completely restored mitochondrial elongation, similar to the effect of expressing MFN2-MYC (Supp Fig. 2A, B). Strikingly, expression of the MFN2-IYFFT in MTCH2 KO cells was sufficient to restore mitochondrial elongation, while expression of MFN2-ACTA had a less strong effect (Fig. 2A, B, C). These results suggest that in terms of mitochondrial morphology, increasing MFN2’s levels at the ER-mitochondrial contact sites is sufficient to compensate for the loss of MTCH2. To test whether MFN2-YIFFT rescue effect could be supplanted by overexpression of an artificial ER-mitochondria tethering protein, we used the artificial linker AKAP-mtagBFP2-UBC6 (Hirabayashi *et al*., 2017), and found no recue of MTCH2 KO mitochondrial fragmentation (Supp Fig. 2C, D). These results suggest that MFN2 possesses a specific function which cannot be supplanted with an artificial ER-mitochondria tether.

**Figure 2.**
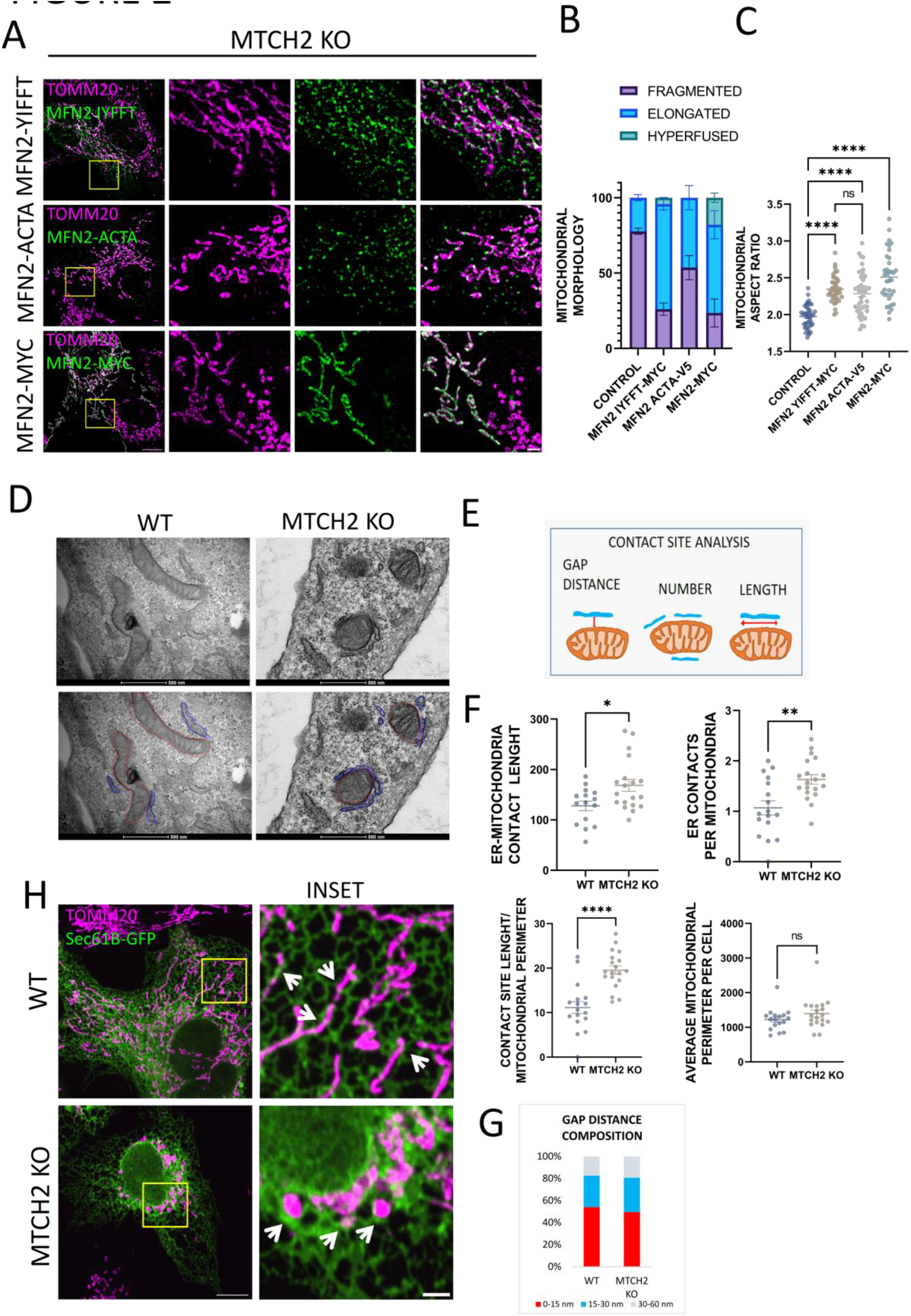
MFN2-targeted to the ER is sufficient to restore mitochondrial elongation in MTCH2 KO cells, and loss of MTCH2 results in increased ER-mitochondria contact sites. **A** Representatives images of MTCH2 KO MEFs transfected with MFN2-IYFFT-MYC, MFN2-ACTA-V5 or MFN2-MYC. Mitochondria were stained with αTOMM20 antibody. Scale bar 10 μm; inset scale bar 2 μm. **B** Classical classification of mitochondrial network morphology. MTCH2 KO non-transfected (control) or transfected with MFN2-IYFFT, MFN2-ACTA or MFN2-MYC. Results are presented as average and SEM of three separate experiments; at least 20 cells were analyzed per condition. **C** Mitochondrial aspect ratio analysis of MTCH2 KO MEFs non-transfected (control) or transfected with MFN2-IYFFT, MFN2-ACTA or MFN2-MYC; at least 20 control or transfected cells were analyzed per condition. One-way ANOVA statistical analysis. **D** Upper panel: Electron micrographs showing representative areas depicting mitochondria-ER contact regions in WT and MTCH2 KO cells. Lower panel: free hand modeling of ER and mitochondria. ER regions that were in a distance of 60 nm or less were considered as contact regions and free-hand delineated in blue. Mitochondrial perimeter was delineated in red. Scale bar 500 nm. **E** Schematic representation of ER-mitochondria contact sites analysis. For every ER-mitochondria contact analyzed the distance and length was quantified. Also, the number of ER-mitochondria contact sites per cell was analyzed and normalized to the total number of mitochondria per cell. Finally, using the total mitochondrial perimeter per cell, and the total length of the ER-mitochondria contacting regions the average of the percentage of mitochondrial perimeter covered by contacts with the ER per cell was calculated. **F** Quantification of ER-mitochondria contact sites: average number of ER-mitochondria contact sites per mitochondria; average length of ER-mitochondria contact sites; average mitochondrial perimeter and percentage of mitochondrial perimeter covered by ER contact sites. At least 15 WT and MTCH2 KO cells were analyzed, statistical analysis was performed by t-test. **G** Quantification of ER-mitochondria contact sites GAP distance composition in WT and MTCH2 KO MEFs. **H** Representative images of WT and MTCH2 KO cells transfected with the ER marker Sec61b-GFP and stained with αTOMM20 antibody. Arrowheads indicate areas of close proximity between ER and mitochondria. MTCH2 KO mitochondria show large regions in close proximity with ER and specially enriched in the perinuclear region. Scale bar 10 μm; inset scale bar 2 μm.

Since ER-localized MFN2 was sufficient to rescue loss of MTCH2, we investigated whether MTCH2 deletion leads to a defect in tethering between ER and mitochondria. To evaluate this, we used transmission electron microscopy (EM) and characterized the interaction between ER and mitochondria in WT and MTCH2 KO MEFs. We analyzed GAP distance between mitochondria and ER, the number of ER contacts per mitochondria and the average length of contacts between the organelles (Fig. 2D, E). Interestingly, MTCH2 KO cells showed an increased number of contact sites between ER and mitochondria and an increase in the contact site length (Fig. 2F). These results translate into an overall increase in the total contact surface between the two organelles. On the other hand, MTCH2 deletion had no impact over the distance that separates both organelles as seen in the GAP distance composition analysis (Fig. 2F, bottom panel), suggesting that MTCH2 does not work as a tethering factor between mitochondria and ER.

To evaluate the macro-morphology of the ER and mitochondrial networks at the light microscopy level, we transfected WT and MTCH2 KO MEFs with the ER marker Sec61b-GFP. ER morphology appeared normal in MTCH2 KO MEFs, and in agreement with our EM data, we could observe large overlapping areas between ER membranes and mitochondria, which were enriched in the peri-nuclear region of the cell (Fig. 2G).

Altogether our results indicate that MFN2-targeted to the ER is sufficient to restore mitochondrial elongation in MTCH2 KO cells, and that MTCH2 KO cells show increased ER-mitochondria contact sites.

### MFN2-mediated mitochondrial fusion requires LPA synthesis

Since MTCH2 deletion increases the number and length of ER-mitochondria contact sites, and MFN2-targeted to the ER restores mitochondria elongation in MTCH2 KO cells, we hypothesized that the ER may play a role in preserving mitochondrial plasticity in MTCH2 KO cells. MFN2’s mitochondrial-ER tethering activity was shown to be necessary for calcium flux (Deborah *et al*., 2016; Gautier *et al*., 2016; Han *et al*., 2021) as well as for lipid transfer (Area-Gomez *et al*., 2012; Hernández-Alvarez *et al*., 2019). Several studies highlight the requirement of MTCH2 for normal mitochondrial lipid biosynthesis (Kulyté *et al*., 2011; Bernhard *et al*., 2013; Bar-Lev *et al*., 2016; Landgraf *et al*., 2016; Rottiers *et al*., 2017). Therefore the lipid species known to promote mitochondrial fusion including lysophosphatidic acid (LPA), phosphatidic acid (PA) and cardiolipin (CL) (Ohba *et al*., 2013; Zhang *et al*., 2016; Kameoka *et al*., 2018), were candidate targets for mediating MTCH2 dependent enforced mitochondrial fusion. Consistent with this idea, it was recently described that MTCH2 funnels the activity of the bioactive LPA towards the mitochondrial fusion machinery (Labbé *et al*., 2021).

In light of the current findings, we evaluated the importance of LPA synthesis for mitochondrial fusion enforced by either MTCH2 or by MFN2. To inhibit LPA synthesis we used the GPAT inhibitor FSG67 (Wydysh *et al*., 2009; Kuhajda *et al*., 2011), which was designed to inhibit GPAT1 but was shown to also inhibit GPAT2 and GPAT3 activity (Kuhajda *et al*., 2011; Clemens *et al*., 2019). First, we evaluated the effect of inhibiting LPA synthesis on mitochondrial morphology in WT MEFs. WT MEFs exposed for 16 hours to FSG67 at a concentration that does not affect cellular viability (McFadden *et al*., 2014), elicited mitochondrial fragmentation with complete penetrance, which was reversible after washing the inhibitor (Fig. 3A, B).

**Figure 3.**
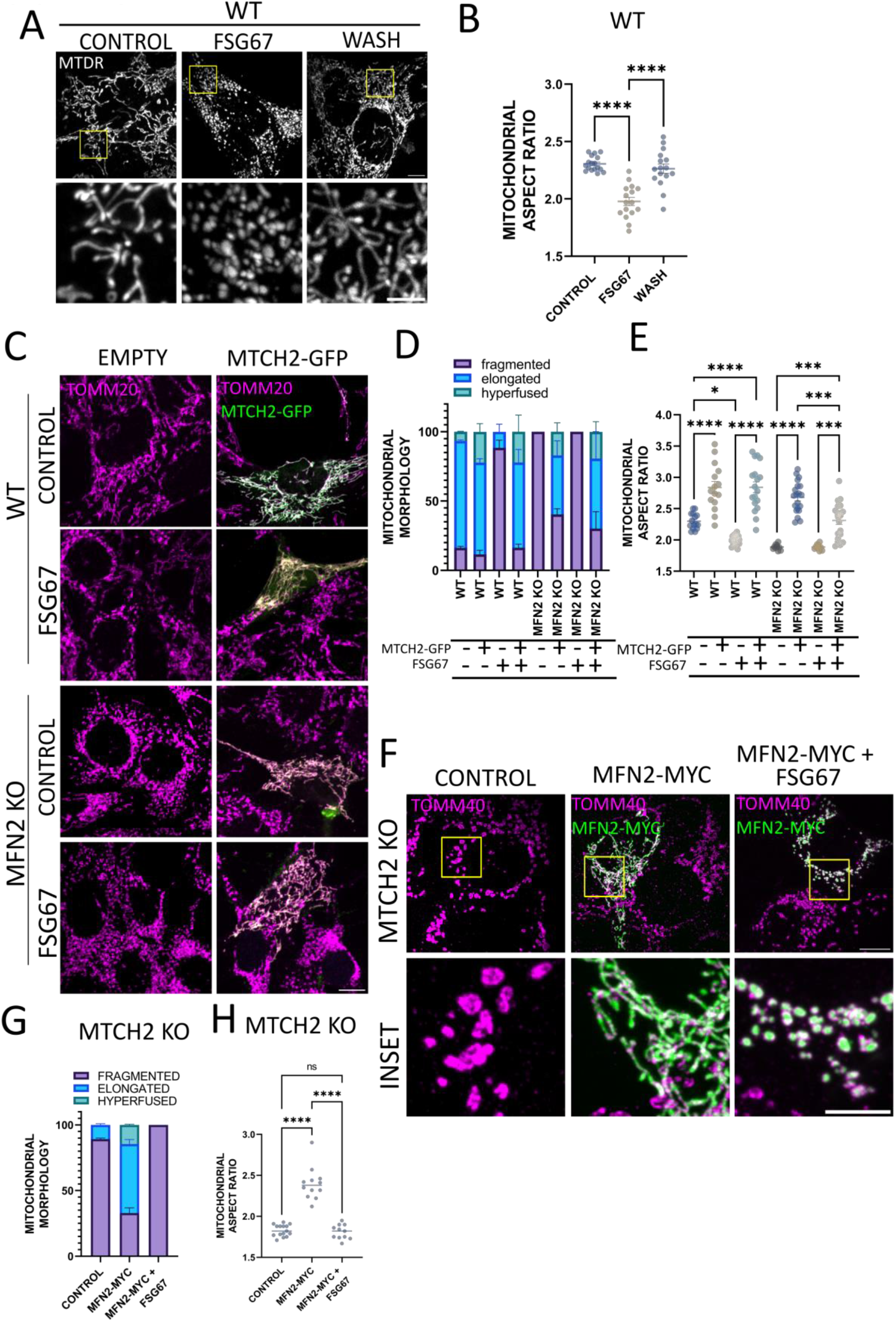
MFN2-mediated mitochondrial fusion requires LPA synthesis. **A** Representatives images of WT MEFs non-treated (control) or treated for 16 hrs with 125 μM FSG67 -/+ wash, and imaged after 4 hrs. Mitochondria were labeled with MTDR. Scale bar 10 μm; Inset scale bar 5 μm. **B** Mitochondrial aspect ratio analysis of cells presented in A. One-way ANOVA statistical analysis. **C** Representatives images of WT and MFN2 KO MEFs transfected with empty vector or overexpressing MTCH2-GFP in control conditions or treated for 16 hrs with FSG67. Mitochondria were stained with αTOMM20 antibody. Scale bar 10 μm. **D** Classical classification of mitochondrial network morphology of WT and MFN2 KO MEFs transfected with empty vector or overexpressing MTCH2-GFP in control conditions or treated for 16 hrs with FSG67. Results are presented as average and SEM of three separate experiments; at least 20 cells were analyzed per condition. **E** Mitochondrial aspect ratio analysis of WT and MFN2 KO MEFs transfected with empty vector or overexpressing MTCH2-GFP in control conditions or treated for 16 hrs with FSG67. At least 15 control and transfected cells were analyzed per condition. One-way ANOVA statistical analysis. **F** Representatives images of MTCH2 KO MEFs transfected with empty vector or overexpressing MFN2-MYC in control conditions or treated for 16 hrs with FSG67. Mitochondria were stained using αTOMM40 antibody. Scale bar 10 μm; Inset scale bar 5 μm. **G** Classical classification of mitochondrial network morphology of MTCH2 KO MEFs transfected with empty vector or overexpressing MFN2-MYC in control conditions or treated for 16 hrs with FSG67. Results presented as average and SEM of three separate experiments; at least 20 cells were analyzed per condition. **H** Mitochondrial aspect ratio analysis of MTCH2 KO MEFs transfected with empty vector or overexpressing MFN2-MYC in control conditions or treated for 16 hrs with FSG67. At least 15 control and transfected cells were analyzed per condition. One-way ANOVA statistical analysis.

If indeed MTCH2 funnels LPA towards the mitochondrial fusion machinery to enable mitochondrial fusion (Labbé *et al*., 2021), then reducing LPA synthesis by inhibiting GPAT activity would likely impair mitochondrial fusion enforced by MTCH2 overexpression. To evaluate this, we transfected WT and MFN2 KO cells with MTCH2-GFP, and treated the cells with FSG67. Unexpectedly, MTCH2-GFP overexpression still elicited mitochondrial elongation in WT cells treated with FSG67 and to a lesser extent in MFN2 KO cells treated with FSG67 (Fig. 3C-E), suggesting that reducing LPA levels by inhibiting GPAT activity is insufficient to impair MTCH2-mediated mitochondrial fusion.

To confirm these results in another cellular system, we knocked out MTCH2 KO in human HEK293T cells using CRISPR/Cas9 gene editing (Supp Fig. 3A). In line with ours and others findings in MEFs, mouse embryonic stem cells (Bahat *et al*., 2018), and HCT116 cells (Labbé *et al*., 2021), MTCH2 deletion in HEK293T cells elicits extensive mitochondrial fragmentation (Supp Fig. 3B, C). Also treatment of WT HEK293T cells with FSG67 elicited mitochondrial fragmentation, which was reversed upon washing the inhibitor (Supp Fig. 3D, E). Similar to the results obtained in MEFs, MTCH2-GFP overexpression elicited mitochondrial tubulation in WT HEK293T cells treated with FSG67 (Supp Fig. 3F, G), further confirming the notion that inhibition of LPA synthesis is insufficient to block MTCH2’s ability to induce mitochondrial fusion.

Next, we evaluated whether inhibiting LPA synthesis would have an effect on MTCH2-independent mitochondrial fusion. For this purpose, we overexpressed MFN2-MYC in MTCH2 KO MEFs, and left the cells either untreated or treated with FSG67. Inhibition of LPA synthesis completely inhibited MFN2’s ability to restore mitochondrial elongation in MTCH2 KO cells (Fig. 3F-H). Similar results were observed in MTCH2 KO HEK293T cells, where MFN2 overexpression elicited mitochondrial elongation in MTCH2 KO cells, and this effect was abolished by FSG657 (Supp Fig. 3F-H). These findings indicate that in the absence of MTCH2, MFN2-induced mitochondrial elongation shows an absolute dependence on LPA synthesis.

### MTCH2 cooperates with MFN2 and lysophosphatidic acid synthesis to sustain mitochondrial fusion under cellular stress conditions

To further understand the interplay between MTCH2, MFN2 and LPA synthesis in mitochondrial fusion, we choose to enforce mitochondrial fusion using different known stresses (Tondera *et al*., 2009; Gomes, Benedetto and Scorrano, 2011; Rambold *et al*., 2011). It was recently proposed that MTCH2 plays a specific role in mediating mitochondrial fusion in response to amino-acid deprivation since HCT116 MTCH2 KO cells starved for amino-acids were unable to elongate their mitochondria (Labbé *et al*., 2021). We found that MTCH2 KO MEFs did not show a similar defect and were able to elongate their mitochondria after amino-acids deprivation, induced by HBSS treatment (Supp Fig. 4A-C). The hyperfusion observed upon amino-acid deprivation was fully dependent on the synthesis of LPA, since pretreatment with FSG67 prior to HBSS-treatment led to cell death in both WT and MTCH2 KO cells (Supp Fig. 4A-C). HBSS treatment had no effect on mitochondrial morphology in WT and MTCH2 KO HEK293T cells (Supp Fig. 4F), suggesting that the mitochondrial response to this stress and the requirement for MTCH2 may vary among different cell lines. Overall, our data suggest that MTCH2 is not essential for HBSS-induced mitochondrial fusion in MEFs, while MEFs treated with a combination of HBSS and FSG67 do not survive, suggesting increased sensitivity to LPA availability during amino-acid deprivation.

**Figure 4.**
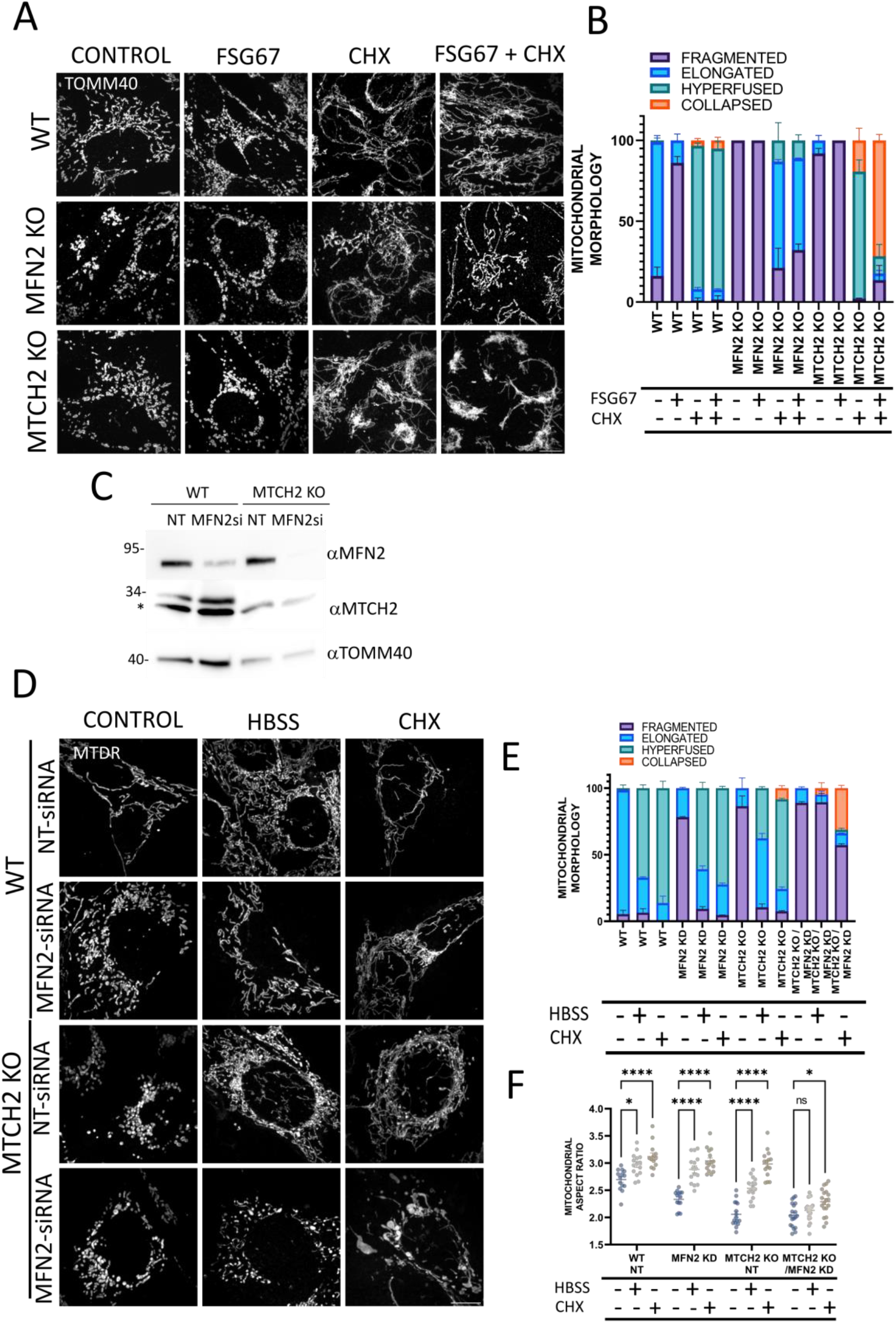
MTCH2 cooperates with MFN2 and lysophosphatidic acid synthesis to sustain mitochondrial fusion under cellular stress conditions. **A** Representatives images of WT, MFN2 KO and MTCH2 KO MEFs either non-treated (control) or treated with FSG67 for 16 hrs, treated with 10 μM CHX for 4 hrs or pretreated for 16 hrs with FSG67 and then treated with 10 μM CHX for 4 hrs in the presence of the inhibitor. Mitochondria were stained with αTOMM40 antibody. Scale bar 10 μm. **B** Classical classification of mitochondrial network morphology of cells presented in A. Results are presented as average and SEM of three different experiments, and at least 30 cells were analyzed per condition. **C** Western blot analysis of MFN2 in WT and MTCH2 KO MEFs treated with non-targeting (NT) siRNA or with siRNA targeting MFN2 expression, and TOMM40 serves as loading control. * Depicts a non-specific band. **D** Representative images of WT and MTCH2 KO MEFs treated with non-targeting (NT) siRNA or with MFN2 siRNA, in control conditions, treated with HBSS for 4 hrs or treated with 10 μM CHX for 4 hrs. Mitochondria were labeled with MTDR. Scale bar 10 μm. **E** Classical classification of mitochondrial network morphology of cells shown in D. Results are presented as average and SEM of three different experiments, and at least 30 cells analyzed per condition. **F** Mitochondrial aspect ratio analysis of cells shown in D. At least 15 cells were analyzed per condition. One-way ANOVA statistical analysis.

Since amino-acid deprivation is lethal when LPA synthesis is inhibited, we induced stress-induced mitochondrial hyperfusion (SIMH) by treating cells with the protein translation inhibitor cycloheximide (CHX) (Tondera *et al*., 2009). In agreement with previous studies (Tondera *et al*., 2009; Labbé *et al*., 2021), WT, MTCH2 KO and MFN2 KO mitochondria fuse in response to SIMH, while MFN1 KO and MFN1/2 DKO mitochondria remain fragmented (Supp Fig. 4D, E). These results suggest that MTCH2 as well as MFN2 are not, whereas MFN1 is, necessary for CHX induced-SIMH, supporting the notion of a more important role for MFN1 over MFN2 in sustaining mitochondrial fusion (Ishihara, Eura and Mihara, 2004).

Next, we evaluated the importance of LPA synthesis for SIMH by pretreating WT, MTCH2 KO and MFN2 KO cells with FSG67 prior to addition of CHX. WT cells showed mitochondria hyperfusion in response to CHX in the presence of FSG67, while MFN2 KO cells showed mitochondria elongation but not mitochondria hyperfusion under the same conditions (Fig. 4A, B). These results suggest that WT and MFN2 KO cells do not require full GPAT activity and newly synthesized LPA for stress-induced mitochondrial elongation/hyperfusion. In contrast, FSG67 largely impaired the response of MTCH2 KO cells to SIMH, eliciting mitochondria clumping, resulting in collapse of the mitochondrial network around the nucleus (Fig. 4A, B). Similar results were obtained with HEK293T cells, in which WT cells treated with FSG67 responded normally to SIMH, while FSG67 treated MTCH2 KO cells are less responsive to SIMH (Supp Fig. 4H, I). These results suggest that MTCH2 KO cells, but not MFN2 KO cells, require newly synthesized LPA to respond to SIMH.

Our results until now suggest the coexistence of two pathways: one independent of MTCH2, which can be compensated by MFN2 overexpression and relies on LPA synthesis, and the other independent of MFN2, which can be compensated by MTCH2 overexpression but does not require full GPAT activity. Thus, we envisioned that if loss of one of these pathways elicits mitochondria to rely on the other pathway to sustain mitochondrial plasticity, then loss of both pathways should largely impair mitochondrial fusion.

To test this idea, we silenced MFN2 in MTCH2 KO cells and exposed them to amino-acid deprivation or to SIMH. Similar to MFN2 KO MEFs, MFN2 silencing resulted in mitochondrial fragmentation and did not impair mitochondrial elongation in response to HBSS treatment or SIMH (Fig. 4D-F). Importantly, both MEFs and HEK293T MTCH2 KO cells, in which MFN2 was silenced, showed aberrant or no mitochondrial plasticity after enforced mitochondrial elongation induced by either HBSS or CHX treatment. Thus, our results suggest that mitochondria require the presence of either MTCH2 or MFN2 to sustain plasticity under stress conditions, and that loss of MTCH2 together with either loss of MFN2 or together with LPA synthesis inhibition, impairs this plasticity.

## Discussion

Our results are consistent with the idea of the existence of two complementary pathways required to sustain mitochondrial plasticity/fusion. Loss of MFN2 unmasks the requirement for MTCH2 to sustain mitochondrial plasticity under cellular stress conditions, while loss of MTCH2 unmasks the requirement for LPA synthesis to drive mitochondrial fusion, a process that is facilitated by MFN2-mediated mitochondrial-ER contacts (Fig. 5).

**Figure 5.**
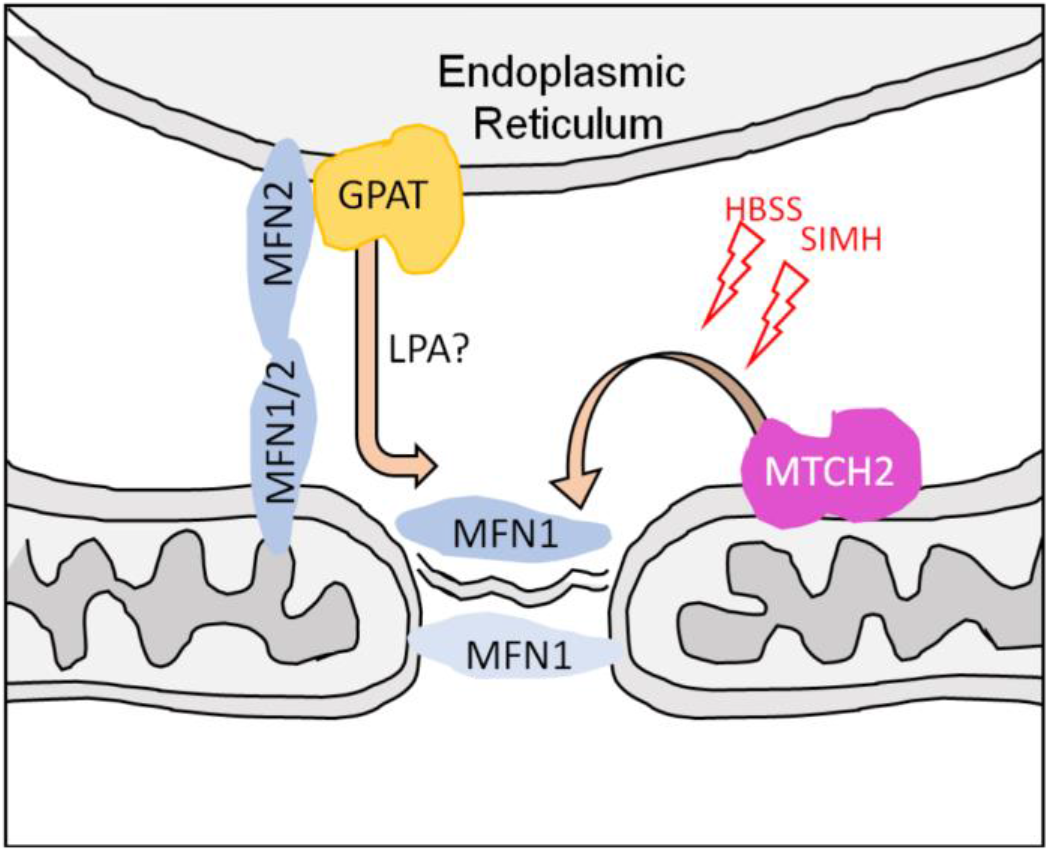
MTCH2 cooperates with MFN2 and LPA synthesis to sustain mitochondrial fusion. MFN2 and GPAT/LPA synthesis (ER) cooperate with MTCH2 (mitochondria) to enable mitochondrial fusion/plasticity induced by cellular stress (HBSS and SIMH).

Based on several examples, it is clear that MFN2 expression is not essential for mitochondrial fusion (Meeusen, McCaffery and Nunnari, 2004; Gomes, Benedetto and Scorrano, 2011; Rambold *et al*., 2011; Palmer *et al*., 2013). Here we demonstrate that MTCH2 also bypasses the requirement of MFN2 for mitochondrial fusion, and that in cells lacking MTCH2, MFN2 becomes essential to sustain mitochondrial plasticity under cellular stress conditions. Thus, we uncovered the contribution of two redundant/cooperative mechanisms that sustain mitochondrial plasticity which are interconnected by the pro-mitochondrial fusion lipid LPA. In addition, using the MTCH2 KO cellular models, we reveal new and important aspects of the tethering role of MFN2 for mitochondrial fusion, and show that loss of MTCH2 unmasks the requirement of de-novo LPA synthesis to sustain mitochondrial fusion and plasticity.

What is the role of MTCH2 in mitochondrial fusion and how does it compensate for MFN2 absence? Recently two different aspects of MTCH2’s function were brought together in a model that suggests that MTCH2 plays a role in funneling LPA towards the mitochondrial fusion machinery (Labbé *et al*., 2021). Our results show that: 1) in cells expressing endogenous MTCH2 (WT and MFN2 KO cells), inhibiting GPAT activity elicits mitochondrial fragmentation, but does not inhibit mitochondrial plasticity; 2) mitochondria of WT and MFN2 KO cells overexpressing MTCH2 do not fragment upon GPAT inhibition, and 3) loss of MTCH2 unmasks the requirement of full GPAT activity and de-novo LPA biosynthesis for mitochondria to retain the ability to respond to fusion-induced by cellular stress.

Since MTCH2 is not essential to sustain mitochondrial fusion, we can infer that MTCH2 funneling activity is not essential for mitochondria to fuse but rather increases its efficiency. Consistent with this idea are the findings showing that in MTCH2 KO cells, mitochondrial fusion events associated with MFN1-GFP foci were rare and the tethering time between mitochondria was long. In addition, MTCH2 deletion has no effect on the assembly of MFN1 and MFN2 native protein complexes. These results suggest that MTCH2 deletion most likely elicits a defect that occurs after docking of mitochondria, and could be connected to inadequate lipidic composition at fusion sites.

Why do MTCH2 and MFN2 complement each other, and enforce mitochondrial fusion independently? Two key observations were important to understand the interplay between MTCH2 and MFN2. First, in MTCH2 KO cells, MFN2’s tethering role is sufficient to enforce mitochondrial fusion and second, inhibiting GPAT activity impairs MFN2’s ability to enforce MTCH2’s-independent mitochondrial fusion. If MTCH2 deletion increases the requirement of LPA to sustain mitochondrial fusion, it is possible that overexpressing MFN2 at ER might be critical to enforce the flux of lipids allowing MTCH2 KO mitochondria to fuse. It was previously shown that MFN2 is necessary for transfer of phosphatidyl ethanolamine from the ER to the mitochondria (Area-Gomez *et al*., 2012; Hernández-Alvarez *et al*., 2019), but so far it has not be shown that MFN2 could also mediate LPA transfer from the ER to the mitochondria. In line with this idea, MTCH2 deletion elicits an increase in the contact area between mitochondria and ER which could reflect an attempt of the mitochondria to compensate for insufficient LPA levels required to sustain mitochondrial fusion. In agreement with this hypothesis, inhibiting LPA synthesis, which impacts GPAT activity in both mitochondria and ER, impairs MFN2-enforced mitochondrial fusion in MTCH2 KO cells, further suggesting that MFN2’s activity required to enforce MTCH2-independent mitochondrial fusion is coupled to lipid biosynthesis and probably to lipid transfer.

The idea that the ER represents a source for pro-mitochondrial fusion lipids has not been explored yet. Based on our observations, it is tempting to speculate that mitochondrial fragmentation in MFN2 KO cells could arise, in part, due to insufficient amounts of pro-mitochondrial fusion lipids coming from the ER, instead of being a consequence of decreased mitochondrial fusion per se. Moreover, MTCH2 overexpression might enforce mitochondrial fusion in MFN2 KO cells by supplementing the absence of LPA and/or other lipids coming from the ER, and funneling them toward mitochondrial fusion sites

Are contact sites and metabolite/lipid transfer actively involved in the mitochondrial fusion event, or are they occurring at different mitochondrial regions in parallel? Two reports provide evidence for the presence of ER contacts during mitochondrial fusion (Guo *et al*., 2018; Abrisch *et al*., 2020), yet what is the functional importance of ER-mitochondrial contact sites in this context has not been addressed yet. It will be important to determine if MFN2’s function as a tethering factor is important to actively incorporate ER contact sites into the mitochondrial fusion event to generate the correct lipidic environment, or whether MTCH2 accumulates at fusion sites prior to mitochondrial fusion events.

It is tempting to propose that MTCH2 regulates the lipidic composition of the OMM to provide a more fusogenic environment, and once inhibited, a pro-fragmentation environment, for situations of cellular stress such as apoptosis. In the future, it would be interesting to investigate how the connection between truncated tBID/full-length BID and MTCH2 might modulate mitochondrial lipid metabolism to elicit mitochondrial fragmentation to enable cytochrome c release at the early stages of apoptosis. And more generally, it will be interesting to address whether MTCH2 might act as a sensor in the OMM that transduces incoming signals into metabolic and architectural adaptations of the mitochondrial network. Our work sheds new light on the complexity of the interaction between mitochondrial dynamics, lipid metabolism and inter-organelle contacts in maintaining mitochondrial architecture and regulating its behavior and plasticity.

## Materials and Methods

### Cell lines and culture conditions

MFN1 KO, MFN2 KO and MFN1/2 DKO MEFs, originally published by Prof. David Chan (Chen *et al*., 2003), were a gift from Prof. Gyorgy Hajnoczky. MEFs and HEK293 cells were generated in our laboratory. MTCH2 KO MEFs were generated from *MTCH2*^*F/F*^ primary MEFs, which were prepared from 11 to 13-day-old embryos and transformed with SV40 (Zaltsman *et al*., 2010). *MTCH2*^*F/F*^ MEFs were transfected with Cre-GFP, and several MTCH2 KO clones were obtained, showing similar mitochondrial fragmentation, and one of these clones was used for this study. The MTCH2 CRISPR KO cell line was generated in HEK293T cells. Guides were designed using CHOP web application (https://chopchop.cbu.uib.no/). MTCH2 guides-RNAs (F-CACCAGCACTTTCACGTACATGAGGT, R-TAAAACCTCATGTACGTGAAAGTGCT). Cells were transfected with guides and selected with puromycin; several MTCH2 KO clones were obtained, showing similar mitochondrial fragmentation, and one of these clones was used for this study. Cells were grown in high glucose DMEM (cat. # 41965, Thermo Fisher), 10% FBS (cat. # 12657, Gibco), supplemented with 2mM L-glutamine (cat. # 03-020, Biological Industries), 1mM sodium pyruvate (cat. # 03-042, Biological Industries) and antibiotics (cat. # 03-031, Biological Industries). For HBSS starvation, cells were washed 3 times with DMEM (cat. # A14430, Thermo Fisher), and then incubated for 4 hrs with HBSS (cat. # 02-015, Biological Industries). CHX 10 μM (cat. # 01810, Milipore-Sigma), and FSG67 125 μM (cat.# 1158383-34-6, Focus Biomolecules) were used in this study.

### Antibodies

Mouse monoclonal antibodies were used for: SDHA (ab14715, Abcam), MYC (SC-40, Santa Cruz), MFN2 (clone 4H8, Sigma), OPA1 (BD612606, BD Transduction Laboratories), Complex III (UQCRC2) (Ab14745, Abcam), V5 tag (R960-25, Invitrogen), GAPDH (sc-47724, Santa Cruz), Tubulin (sc-5286, Santa Cruz), and Complex V (ATP5b) (MS-503, MitoScience). Rabbit polyclonal antibodies were used for: MTCH2 (Ab174921, Abcam), TOMM20 (SC-11418, Santa Cruz), TOMM40 (18409-AP, Proteintech) and MYC (2278, Cell Signaling). MFN1 polyclonal antibody was a kind gift from Prof. Richard Youle. Anti-MTCH2 polyclonal antibodies that recognize the mouse protein were generated in-house (Grinberg *et al*., 2005). Secondary antibodies donkey anti-mouse or donkey anti-rabbit coupled to Cy2, Cy3 or Cy5 dyes were purchased from Molecular Probes, In., and secondary antibodies HRP-conjugated donkey anti-mouse, donkey anti-rat and donkey anti-rabbit were purchased from Amersham GE.

### Expression plasmids, transfections and gene silencing

The MTCH2-GFP plasmid was generated in our laboratory and was previously described (Grinberg *et al*., 2005). We also generated a N-terminally EGFP-tagged MFN1 plasmid (MFN1-GFP) by cloning the full length ORF of mouse MFN1 (accession number AY174062) into a pEGFP-C1 expression vector (Clonetech). MFN2-MYC (cat. # 23213), AKAP-mtagBFP2-UBC6 (cat. # 105007), Sec61b-GFP (cat. # 121159) were purchased from Addgene. MFN2-IYFFT-MYC and MFN2-ACTA-V5 were a kind gift of Prof. Luca Scorrano. Cells were transfected using the JET-PRIME (MEFs) and polyJET (HEK293T) transfection reagents according to the manufacturer’s instructions (MTI; Global Systems). Most experiments were carried out either 24 or 48 hrs post transfection. On-TARGET plus Smart Pool (Dharmacon) siRNAs were used: MFN2 human (L-012961-00), MFN2 mouse (L-046303-00) and non-targeting (NT) control (D-001810-10). Gene silencing was performed by transfecting cells using Lipofectamine RNAi MAX (Invitrogen, 13778150), according to the manufacturer’s instructions, and 72 hrs post transfection gene silencing was confirmed by Western blot analysis.

### Sample preparation and Western blot analysis

Whole cell lysates were obtained by lysing cell pellets with RIPA buffer for 15 minutes on ice. Then samples were centrifugated, and protein concentration was estimated. Subcellular fractionation was performed as described previously (Zaltsman *et al*., 2010). Briefly, cells were collected by scraping, centrifuged and washed with ice cold PBS. For mitochondrial purification, cells were resuspended in mitochondrial buffer (MB, 200 mM mannitol, 70 mM sucrose, 1 mM EGTA, 10 mM HEPES pH 7.5). Cells were broken using a 27.5-gauge syringe, and cell debris and unbroken cells were pelleted at low-speed centrifugation (900 g, 10 minutes, 4°C). The supernatant was then centrifuged at high speed (10,000 g, 10 minutes, 4°C) to obtain the heavy membrane fraction enriched in mitochondria. Protein concentration was assessed by Bradford, and equal amounts of protein were separated by SDS-PAGE. Size separated denaturized samples were then transferred to either PVDF or nitrocellulose membranes, blocked in milk, incubated with different primary antibodies followed by peroxidase-conjugated anti-rat, anti-mouse or anti-rabbit, followed by ECL treatment (Amersham).

### Blue Native Gel Electrophoresis (BNGE)

For isolation of native mitochondrial protein complexes we proceeded as previously described (Gomes *et al*., 2011), with small modifications. Briefly, 200 μg of mitochondria were resuspended in 50 μl of native loading buffer (Invitrogen) containing 4% digitonin (Sigma) and protease-inhibitor cocktail (Sigma). After incubating 10 min at 4°C, the lysate was cleared by centrifugation at 22000 *g* for 30 min at 4°C. 2.5 μl of native additive G250 5% (Invitrogen) was added to the supernatant and 100 μg of protein was loaded onto a 3-12% native gel (NativePAGE™ (Invitrogen)). Native complexes were size separated for 30 min at 150V with Dark Blue cathode buffer and 90 min at 250V with Light Blue cathode buffer (Invitrogen). Finally, the native protein complexes were transferred to a PVDF membrane using Native Transfer (Novex) and Western blotted with different antibodies.

### Microscopy studies

Confocal images were acquired in a Nikon ECLIPSE Ti2-E inverted microscope with CSW-1 spinning disc system (Yokogawa), with a ×100 CFI Plan Apo100x oil (na 1.45 wd 0.13mm), equipped with temperature and CO_2_ control. Images were acquired using a Photometrics prime 95b camera and Nikon Elements software. Lasers 405/488/561/638 nm were used for DAPI/BFP, GFP, Cy3 and Cy5/MTDR respectively. Live imaging time lapse experiments were acquired in a wide field Olympus IX83 microscope (Olympus, Japan) equipped with ×100 oil immersion objective Plan Apo100x oil (na 1.45 wd 0.13mm), Orca Flash 4.0 camera (Hamamatsu Photonics, Japan) and a LED light source (CoolLED, UK). Images were acquired using cellSense software and preprocessed using the online deblur tool.

### Immunofluorescence

For immunofluorescence studies, cells were seeded 24 hrs before the experiment in 13 mm sterile glass coverslips (Thermo scientific). Cells were then washed with PBS and fixed with 4% PFA for 15 min at 37°C. PFA was quenched by incubating the samples for 10 min in 50 mM NH_4_Cl_2_ in PBS. After washing, cells were permeabilized with 0.1% triton x-100 in PBS for 20 min and then blocked for 1 hr at room temperature with 5% BSA, 0.1% triton x-100 in PBS. Cells were then probed with primary antibodies, washed and stained with secondary antibodies conjugated to Cy2, Cy3 or Cy5 (Molecular Probes, Inc.). After three additional washes with DDW, coverslips were mounted on slides using SlowFade Light Antifade kit (Molecular Probes, Inc.).

### Live imaging

For live cell imaging experiments, cells were seeded 24 hrs before the experiment. Cells were pre-incubated with 100 nM Mitotracker Deep Red (MTDR) (Thermo Fisher M22426) for 30 min. Prior to imaging, cells were washed and media was changed to imaging media (DMEM without phenol red, 10% FBS, 2mM glutamine, 1mM sodium pyruvate and 20 mM HEPEs) and stabilized for an additional 30 min at 37°C and 5% CO_2_. Cells were then imaged under temperature and CO_2_-controlled conditions.

### Classical classification of mitochondrial network morphology

Cells were classified according to the morphology of the mitochondrial network divided into three categories according to Karbowski *et al*., 2006: “Elongated” with >90% of mitochondria forming elongated interconnected networks, with >90% round mitochondria. We included the category “Hyperfused” for mitochondrial network highly reticular and interconnected, and “Collapsed” for mitochondrial network that shows clumping surrounding the perinuclear region of the cell. The data presented is an average and standard error of the percentages of three different experiments

### Mitochondrial aspect ratio (AR)

To perform mitochondria AR analysis, images acquired under the same conditions were processed using background subtraction, thresholded (binarized) and then segmented using the Analyze Particles tool from FIJI (Schindelin *et al*., 2012). One representative experiment out of three replicas was analyzed by calculating average mitochondrial AR per cell or average mitochondrial AR per field of view. At least 15 cells or fields were analyzed in each condition. Statistical analysis was performed by one way ANOVA with multiple comparisons.

### MFN1-mediated mitochondrial tethering time

WT and MTCH2 KO cells were transfected with MFN1-GFP and stained with MTDR, and several cells (approximately 30 fields) were imaged for a period of 5 min, taking one image every 3 sec. We located and isolated 23 WT and 18 MTCH2 KO different MFN1-GFP tethering events, which we defined as those events where mitochondria remained visibly tethered through an MFN1-GFP foci for more than 15 seconds, trying to avoid “kiss and run” type of interactions (Liu *et al*., 2009). We utilized the Particle Tracking of the MOSAIC plugin on FIJI to track MFN1-GFP foci during the fusion attempts, and we measured the time that the MFN1 foci spent in these tethering points.

### EM sample preparation and contact site analysis

For EM sample preparation, we seeded cells 24 hrs prior to collection. Cells were washed and fixed for 1 hr at RT in 0.1 M CaCO buffer (4% PFA, 0.1% glutaraldehyde). Next, the fixative was washed using 0.1 M CaCO buffer and incubated O/N in CaCO buffer at 4°C. Next, cells were carefully scrapped, and pelleted. Cell pellets were dehydrated and included in epoxy resin. Finally, samples were cut in 70-micron slices and stained with heavy metals. To analyze ER-mitochondria contact sites by EM, we used the freehand drawing tool from FIJI software. We delimited only ER tubules within a 60 nm distance from the nearest mitochondrial surface, and delimited the mitochondrial perimeter. Then, we analyzed length and GAP of each contact site. We separated the contact site according to their GAP into three groups: from 0-15 nm, 15-30 nm and 30-60 nm. Finally, we calculated the amount of contacts per mitochondria, the average contact length, mitochondrial perimeter, total mitochondrial number and ER-mitochondria contact coverage.

## Statistical analysis

P values were calculated by one-way ANOVA test, unless otherwise specified, with GraphPad prism statistical software.

## Acknowledgements

We are grateful to all the members of the Gross laboratory for their support, insightful discussions and comments on the manuscript.

## Author contributions

A. Goldman performed most of the experiments presented in the paper. Y.Z., M.M. helped with many of the experiments described. A. Goldman and A. Gross planned the project and wrote the manuscript. All authors discussed the results and commented on the manuscript.

## Conflict of interest

The authors declare no conflict interests.

## Figures

**Supplementary Figure 1.**
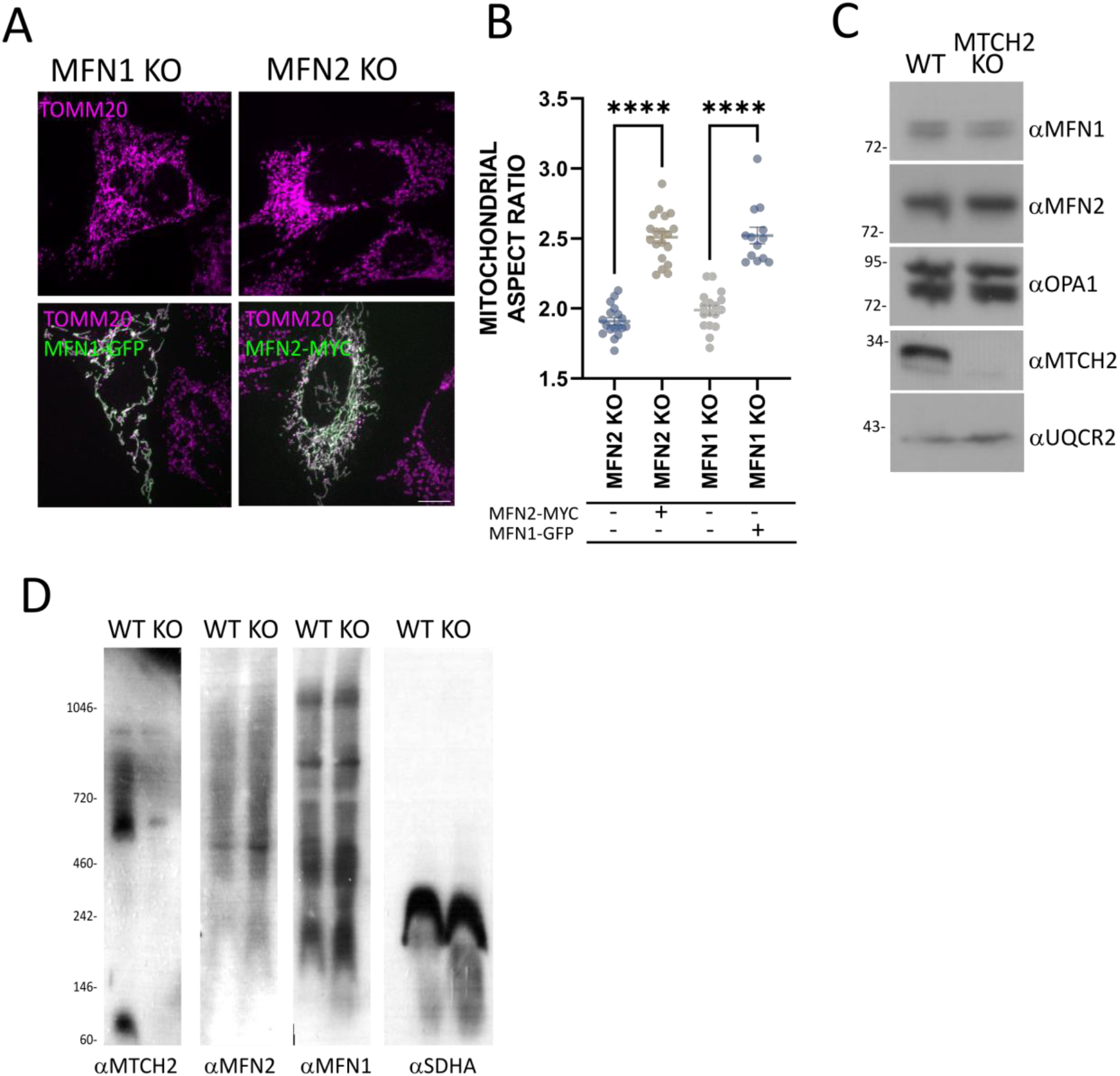
**A** Representative images of MFN1 KO and MFN2 KO MEFs expressing MFN1-GFP or MFN2-MYC, respectively. Mitochondria were stained with αTOMM20 antibody. Scale bar 10 μm. **B** Mitochondrial aspect ratio analysis. At least 20 control and transfected cells were analyzed per condition. One-way ANOVA statistical analysis. **C** Western blot analysis of the mitochondrial fusion proteins MFN1, MFN2 and OPA1 in heavy membrane fractions prepared from WT and MTCH2 KO cells; UQCR2 (complex III) serves as loading control. **D** BN-PAGE of WT and MTCH2 KO heavy membrane fractions blotted with anti-MTCH2, MFN1 and MFN2 antibodies. SDHA serves as control.

**Supplementary Figure 2.**
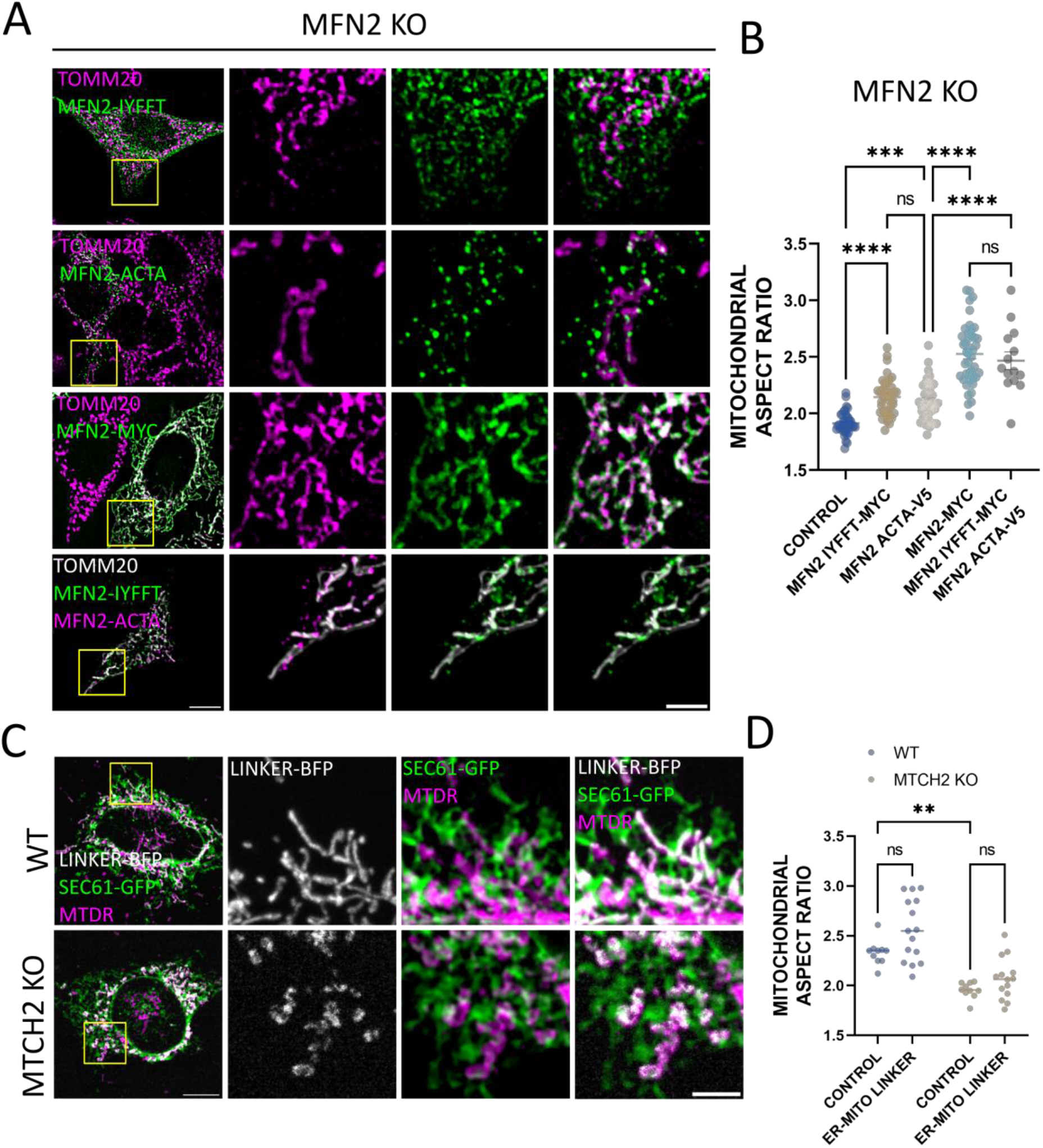
**A** Representatives images of MFN2 KO MEFs transfected with MFN2-IYFFT-MYC, MFN2-ACTA-V5, MFN2-MYC, or co-transfected with MFN2-IYFFT-MYC and MFN2-ACTA-V5. Mitochondria were stained with αTOMM20 antibody. Scale bar 10 μm; inset scale bar 5 μm. **B** Mitochondrial aspect ratio analysis of MFN2 KO MEFs non-transfected (control) or transfected with MFN2-IYFFT, MFN2-ACTA, MFN2-MYC, or co-transfected with MFN2-IYFFT-MYC and MFN2-ACTA. At least 15 control and transfected cells were analyzed per condition. One-way ANOVA statistical analysis. **C** Representatives images of WT and MTCH2 KO MEFs co-transfected with the ER-mitochondria linker construct AKAP-mtagBFP2-UBC6 and the ER marker Sec61b-GFP. Mitochondria were labeled with MTDR. Scale bar 10 μm; Scale bar inset 5 μm. **D** Mitochondrial aspect ratio analysis of WT and MTCH2 KO MEFs non-transfected (control) or transfected with the ER-mitochondrial linker AKAP-mtagBFP2-UBC6. At least 15 control and transfected cells were analyzed per condition. One-way ANOVA statistical analysis.

**Supplementary Figure 3.**
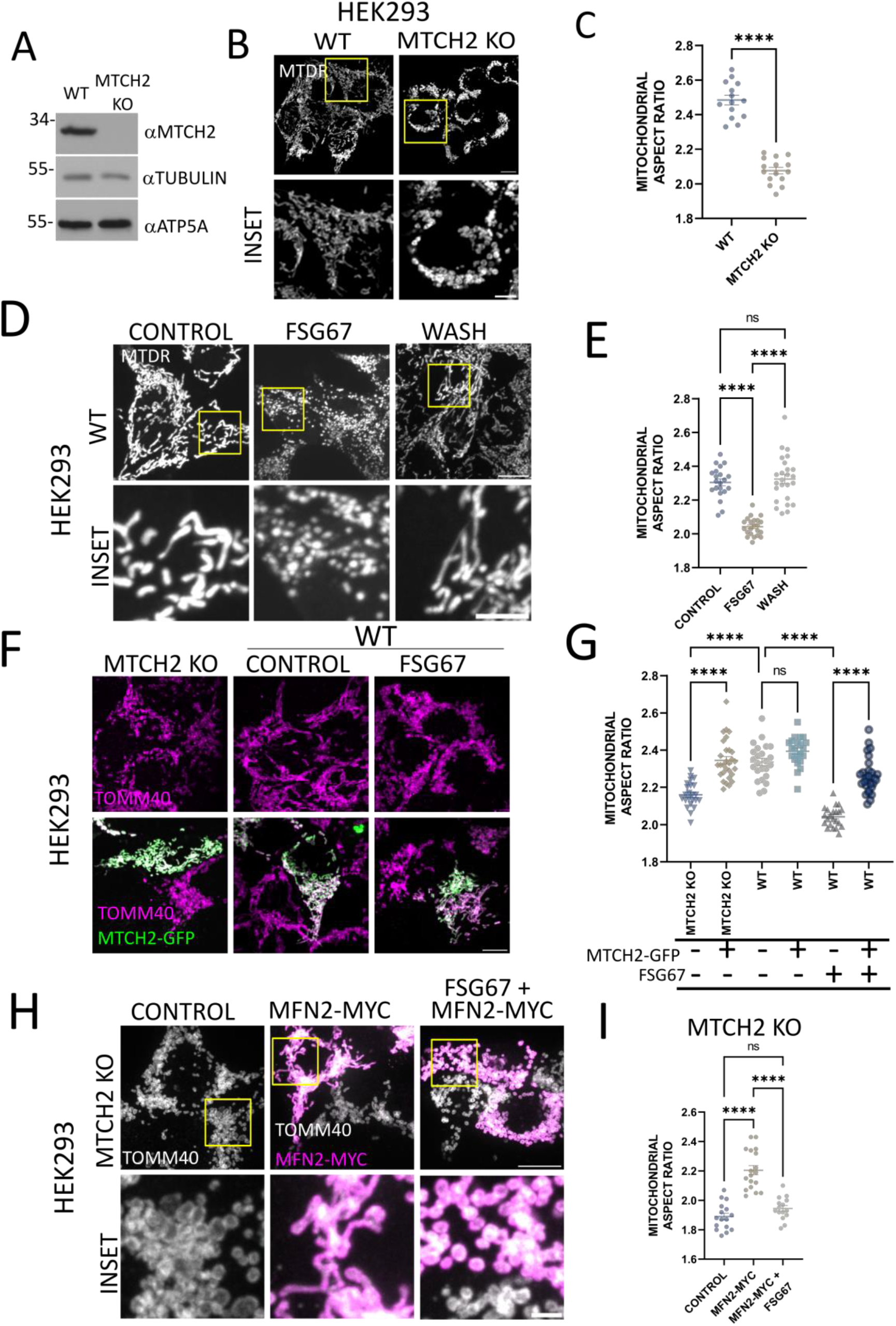
**A** Western blot analysis of MTCH2 in WT and MTCH2 KO HEK293T cells; tubulin and ATPVb (complex V) serve as controls. **B** Representatives images of mitochondria of WT and MTCH2 KO HEK293T cells. Mitochondria were labeled with MTDR. Scale bar 10 μm; Inset scale bar 5 μm. **C** Mitochondrial aspect ratio analysis of WT and MTCH2 KO HEK293T cells. At least 15 fields were analyzed per condition. statistical analysis was performed by t-test. **D** Representatives images of WT HEK293T cells either non-treated (control) or treated for 16 hrs with FSG67 -/+ wash and imaged after 4 hrs. Mitochondria were labeled with MTDR. Scale bar 10 μm; Inset scale bar 5 μm. **E** Mitochondrial aspect ratio analysis of cells presented in D. At least 15 control and transfected cells were analyzed per condition. One-way ANOVA statistical analysis. **F** Representatives images of WT and MTCH2 KO HEK293T cells overexpressing MTCH2-GFP in control conditions and of WT cells treated for 16 hrs with FSG67 and overexpressing MTCH2-GFP. Mitochondria were stained using αTOMM20 antibody. Scale bar 10 μm. **G** Mitochondrial aspect ratio analysis of cells presented in F. At least 15 control and transfected cells were analyzed per condition. One-way ANOVA statistical analysis. **H** Representatives images of MTCH2 KO HEK293T cells transfected with empty vector or overexpressing MFN2-MYC in control conditions or treated for 16 hrs with FSG67. Mitochondria were stained using αTOMM40 antibody. Scale bar 10 μm; Inset scale bar 5 μm. **I** Mitochondrial aspect ratio analysis of cells shown in H. At least 15 control and transfected cells were analyzed per condition. One-way ANOVA statistical analysis.

**Supplementary Figure 4.**
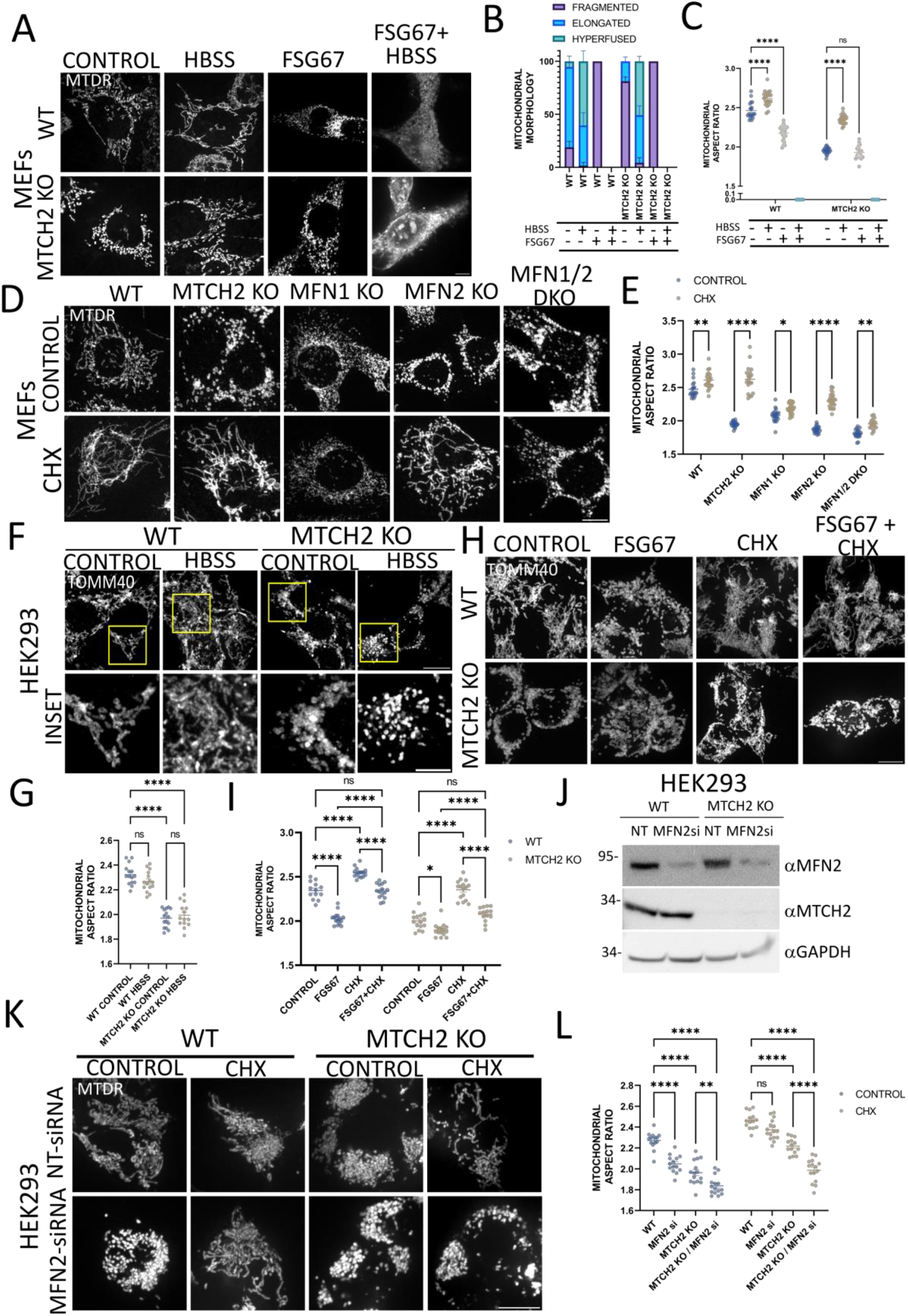
**A** Representatives images of WT and MTCH2 KO MEFs either non-treated (control), treated with HBSS for 4 hrs, treated with FSG67 for 16 hrs, or pretreated for 16 hrs with FSG67 and then treated with HBSS for 4 hours in the presence of the inhibitor. Mitochondria were labeled with MTDR. Scale bar 10 μm. **B** Classical classification of mitochondrial network morphology of cells shown in A. Results are presented as average and SEM of three different experiments, and at least 30 cells analyzed per condition. **C** Mitochondrial aspect ratio analysis of cells shown in A. At least 15 were analyzed per condition. One-way ANOVA statistical analysis. **D** Representatives images of WT and MTCH2 KO, MFN1 KO, MFN2 KO and MFN1/2 DKO MEFs either non-treated (control) or treated with CHX for 4 hrs. Scale bar 10 μm. **E** Mitochondrial aspect ratio analysis of cells shown in D.. At least 15 were analyzed per condition. One-way ANOVA statistical analysis. **F** Representatives images of WT and MTCH2 KO HEK293T cells non-treated (control) or treated with HBSS for 4 hrs. Mitochondria were stained with αTOMM40 antibody. Scale bar 10 μm; Inset scale bar 5 μm. **G** Mitochondrial aspect ratio analysis of cells shown in F. At least 15 were analyzed per condition. One-way ANOVA statistical analysis. **H** Representatives images of WT and MTCH2 KO HEK293T cells either non-treated (control), treated with FSG67 for 16 hrs, treated with CHX for 4 hrs or pretreated for 16 hrs with FSG67 and then treated with CHX for 4 hrs in the presence of the inhibitor. Scale bar 10 μm. **I** Mitochondrial aspect ratio analysis of cells shown in H. At least 15 were analyzed per condition. One-way ANOVA statistical analysis. **J** Western blot analysis of MFN2 in WT and MTCH2 KO HEK293T cells treated with non-targeting (NT) siRNA or with MFN2 siRNA. GAPDH serves as a loading control. **K** Representative images of WT and MTCH2 KO HEK293T cells treated with non-targeting (NT) siRNA or with MFN2 siRNA, in control conditions or treated with CHX for 4 hrs. Mitochondria were labeled with MTDR. Scale bar 10 μm. **L** Mitochondrial aspect ratio analysis of cells shown in K. At least 15 were analyzed per condition. One-way ANOVA statistical analysis.

## Notes

### Competing Interest Statement

The authors have declared no competing interest.

## References

Abrisch, R.G. et al. (2020) ‘Fission and fusion machineries converge at ER contact sites to regulate mitochondrial morphology’, Journal of Cell Biology, 219(4). doi:10.1083/jcb.201911122.

Annis, M.G. et al. (2001) ‘Endoplasmic reticulum localized Bcl-2 prevents apoptosis when redistribution of cytochrome c is a late event’, Oncogene, 20(16), pp. 1939–1952. doi:10.1038/sj.onc.1204288.

Area-Gomez, E. et al. (2012) ‘Upregulated function of mitochondria-associated ER membranes in Alzheimer disease’, The EMBO Journal, 31(21), pp. 4106–4123. doi:https://doi.org/10.1038/emboj.2012.202.

Bahat, A. et al. (2018) ‘MTCH2-mediated mitochondrial fusion drives exit from naïve pluripotency in embryonic stem cells’, Nature Communications, 9(1), p. 5132. doi:10.1038/s41467-018-07519-w.

Bar-Lev, Y. et al. (2016) ‘Mimp/Mtch2, an Obesity Susceptibility Gene, Induces Alteration of Fatty Acid Metabolism in Transgenic Mice.’, PloS one, 11(6), p. e0157850. doi:10.1371/journal.pone.0157850.

Bereiter-Hahn, J. and Vöth, M. (1994) ‘Dynamics of mitochondria in living cells: shape changes, dislocations, fusion, and fission of mitochondria.’, Microscopy research and technique, 27(3), pp. 198–219. doi:10.1002/jemt.1070270303.

Bernhard, F. et al. (2013) ‘Functional relevance of genes implicated by obesity genome-wide association study signals for human adipocyte biology.’, Diabetologia, 56(2), pp. 311–322. doi:10.1007/s00125-012-2773-0.

de Brito, O.M. and Scorrano, L. (2008) ‘Mitofusin 2 tethers endoplasmic reticulum to mitochondria’, Nature, 456, p. 605.

Buzaglo-Azriel, L. et al. (2016) ‘Loss of Muscle MTCH2 Increases Whole-Body Energy Utilization and Protects from Diet-Induced Obesity’, Cell Reports, 14(7), pp. 1602–1610. doi:10.1016/j.celrep.2016.01.046.

Chen, H. et al. (2003) ‘Mitofusins Mfn1 and Mfn2 coordinately regulate mitochondrial fusion and are essential for embryonic development’, The Journal of Cell Biology, 160(2), pp. 189– 200. doi:10.1083/jcb.200211046.

Cipolat, S. et al. (2004) ‘OPA1 requires mitofusin 1 to promote mitochondrial fusion’, Proceedings of the National Academy of Sciences of the United States of America. 2004/10/27, 101(45), pp. 15927–15932. doi:10.1073/pnas.0407043101.

Clemens, M.M. et al. (2019) ‘The inhibitor of glycerol 3-phosphate acyltransferase FSG67 blunts liver regeneration after acetaminophen overdose by altering GSK3β and Wnt/β-catenin signaling.’, Food and chemical toxicology : an international journal published for the British Industrial Biological Research Association, 125, pp. 279–288. doi:10.1016/j.fct.2019.01.014.

Coleman, R.A. and Lee, D.P. (2004) ‘Enzymes of triacylglycerol synthesis and their regulation’, Progress in Lipid Research, 43(2), pp. 134–176. doi:https://doi.org/10.1016/S0163-7827(03)00051-1.

Das, R. and Chakrabarti, O. (2020) ‘Mitochondrial hyperfusion: a friend or a foe.’, Biochemical Society transactions, 48(2), pp. 631–644. doi:10.1042/BST20190987.

Deborah, N. et al. (2016) ‘Critical reappraisal confirms that Mitofusin 2 is an endoplasmic reticulum–mitochondria tether’, Proceedings of the National Academy of Sciences, 113(40), pp. 11249–11254. doi:10.1073/pnas.1606786113.

Gautier, C.A. et al. (2016) ‘The endoplasmic reticulum-mitochondria interface is perturbed in PARK2 knockout mice and patients with PARK2 mutations.’, Human molecular genetics, 25(14), pp. 2972–2984. doi:10.1093/hmg/ddw148.

Gomes, L.C., Di Benedetto, G. and Scorrano, L. (2011) ‘During autophagy mitochondria elongate, are spared from degradation and sustain cell viability.’, Nature cell biology, 13(5), pp. 589–598. doi:10.1038/ncb2220.

Gomes, L.C., Benedetto, G. Di and Scorrano, L. (2011) ‘During autophagy mitochondria elongate, are spared from degradation and sustain cell viability’, Nature Cell Biology, 13, p. 589.

Grinberg, M. et al. (2005) ‘Mitochondrial carrier homolog 2 is a target of tBID in cells signaled to die by tumor necrosis factor alpha.’, Molecular and Cellular Biology, 25(11), pp. 4579–4590. doi:10.1128/MCB.25.11.4579.

Guo, Y. et al. (2018) ‘Visualizing Intracellular Organelle and Cytoskeletal Interactions at Nanoscale Resolution on Millisecond Timescales’, Cell, 175(5), pp. 1430-1442.e17. doi:https://doi.org/10.1016/j.cell.2018.09.057.

Han, S. et al. (2021) ‘The role of Mfn2 in the structure and function of endoplasmic reticulum-mitochondrial tethering in vivo’, Journal of Cell Science, 134(13), p. jcs253443. doi:10.1242/jcs.253443.

Hernández-Alvarez, M.I. et al. (2019) ‘Deficient Endoplasmic Reticulum-Mitochondrial Phosphatidylserine Transfer Causes Liver Disease’, Cell, 177(4), pp. 881-895.e17. doi:https://doi.org/10.1016/j.cell.2019.04.010.

Hirabayashi, Y. et al. (2017) ‘ER-mitochondria tethering by PDZD8 regulates Ca(2+) dynamics in mammalian neurons.’, Science (New York, N.Y.), 358(6363), pp. 623–630. doi:10.1126/science.aan6009.

Ishihara, N., Eura, Y. and Mihara, K. (2004) ‘Mitofusin 1 and 2 play distinct roles in mitochondrial fusion reactions via GTPase activity’, Journal of Cell Science, 117(26), pp. 6535 LP–6546.

Jingsong, C. et al. (2006) ‘Molecular identification of microsomal acyl-CoA:glycerol-3-phosphate acyltransferase, a key enzyme in de novo triacylglycerol synthesis’, Proceedings of the National Academy of Sciences, 103(52), pp. 19695–19700. doi:10.1073/pnas.0609140103.

Kameoka, S. et al. (2018) ‘Phosphatidic Acid and Cardiolipin Coordinate Mitochondrial Dynamics.’, Trends in cell biology, 28(1), pp. 67–76. doi:10.1016/j.tcb.2017.08.011.

Karbowski, M. et al. (2006) ‘Role of Bax and Bak in mitochondrial morphogenesis’, Nature, 443(7112), pp. 658–662.

Kennedy, E.P. (1961) ‘Biosynthesis of complex lipids.’, Federation proceedings, 20, pp. 934–940.

Kuhajda, F.P. et al. (2011) ‘Pharmacological glycerol-3-phosphate acyltransferase inhibition decreases food intake and adiposity and increases insulin sensitivity in diet-induced obesity.’, American journal of physiology. Regulatory, integrative and comparative physiology, 301(1), pp. R116–30. doi:10.1152/ajpregu.00147.2011.

Kulyté, A. et al. (2011) ‘MTCH2 in human white adipose tissue and obesity.’, The Journal of clinical endocrinology and metabolism, 96(10), pp. E1661–5. doi:10.1210/jc.2010-3050.

Labbé, K. et al. (2021) ‘The modified mitochondrial outer membrane carrier MTCH2 links mitochondrial fusion to lipogenesis’, Journal of Cell Biology, 220(11), p. e202103122. doi:10.1083/jcb.202103122.

Landgraf, K. et al. (2016) ‘Loss of mtch2 function impairs early development of liver, intestine and visceral adipocytes in zebrafish larvae.’, FEBS letters. England, pp. 2852–2861. doi:10.1002/1873-3468.12330.

Lee, J.-Y. et al. (2014) ‘MFN1 deacetylation activates adaptive mitochondrial fusion and protects metabolically challenged mitochondria’, Journal of Cell Science, 127(22), pp. 4954 LP–4963.

Lewin, T.M. et al. (2004) ‘Identification of a New Glycerol-3-phosphate Acyltransferase Isoenzyme, mtGPAT2, in Mitochondria *’, Journal of Biological Chemistry, 279(14), pp. 13488–13495. doi:10.1074/jbc.M314032200.

Liu, X. et al. (2009) ‘Mitochondrial “kiss-and-run”: interplay between mitochondrial motility and fusion-fission dynamics.’, The EMBO journal, 28(20), pp. 3074–3089. doi:10.1038/emboj.2009.255.

Maryanovich, M. et al. (2015) ‘An MTCH2 pathway repressing mitochondria metabolism regulates haematopoietic stem cell fate’, Nature Communications, 6. doi:10.1038/ncomms8901.

McFadden, J.W. et al. (2014) ‘Increasing Fatty Acid Oxidation Remodels the Hypothalamic Neurometabolome to Mitigate Stress and Inflammation’, PLOS ONE, 9(12), p. e115642. Available at: https://doi.org/10.1371/journal.pone.0115642.

Meeusen, S., McCaffery, J.M. and Nunnari, J. (2004) ‘Mitochondrial Fusion Intermediates Revealed in Vitro’, Science, 305(5691), pp. 1747 LP–1752. doi:10.1126/science.1100612.

Nunnari, J. et al. (1997) ‘Mitochondrial transmission during mating in Saccharomyces cerevisiae is determined by mitochondrial fusion and fission and the intramitochondrial segregation of mitochondrial DNA.’, Molecular Biology of the Cell, 8(7), pp. 1233–1242.

Ohba, Y. et al. (2013) ‘Mitochondria-type GPAT is required for mitochondrial fusion.’, The EMBO journal, 32(9), pp. 1265–1279. doi:10.1038/emboj.2013.77.

Olichon, A. et al. (2002) ‘The human dynamin-related protein OPA1 is anchored to the mitochondrial inner membrane facing the inter-membrane space’, FEBS Letters, 523(1–3), pp. 171–176. doi:http://dx.doi.org/10.1016/S0014-5793(02)02985-X.

Olichon, A. et al. (2003) ‘Loss of OPA1 perturbates the mitochondrial inner membrane structure and integrity, leading to cytochrome c release and apoptosis’, Journal of Biological Chemistry, 278(10), pp. 7743–7746. doi:10.1074/jbc.C200677200.

Palmer, C.S. et al. (2013) ‘Adaptor Proteins MiD49 and MiD51 Can Act Independently of Mff and Fis1 in Drp1 Recruitment and Are Specific for Mitochondrial Fission’, Journal of Biological Chemistry, 288(38), pp. 27584–27593. doi:10.1074/jbc.M113.479873.

Pellon-Maison, M. et al. (2007) ‘Mitochondrial glycerol-3-P acyltransferase 1 is most active in outer mitochondrial membrane but not in mitochondrial associated vesicles (MAV)’, Biochimica et Biophysica Acta (BBA) - Molecular and Cell Biology of Lipids, 1771(7), pp. 830–838. doi:https://doi.org/10.1016/j.bbalip.2007.04.001.

Rambold, A.S. et al. (2011) ‘Tubular network formation protects mitochondria from autophagosomal degradation during nutrient starvation.’, Proceedings of the National Academy of Sciences of the United States of America, 108(25), pp. 10190–10195. doi:10.1073/pnas.1107402108.

Robinson, A.J., Kunji, E.R.S. and Gross, A. (2012) ‘Mitochondrial carrier homolog 2 (MTCH2): The recruitment and evolution of a mitochondrial carrier protein to a critical player in apoptosis’, Experimental Cell Research, 318(11), pp. 1316–1323. doi:10.1016/J.YEXCR.2012.01.026.

Rojo, M. et al. (2002) ‘Membrane topology and mitochondrial targeting of mitofusins, ubiquitous mammalian homologs of the transmembrane GTPase Fzo’, Journal of Cell Science, 115(8), pp. 1663–1674. doi:10.1242/jcs.115.8.1663.

Rottiers, V. et al. (2017) ‘MTCH2 is a conserved regulator of lipid homeostasis.’, Obesity (Silver Spring, Md.), 25(3), pp. 616–625. doi:10.1002/oby.21751.

Rowland, A.A. and Voeltz, G.K. (2012) ‘Endoplasmic reticulum–mitochondria contacts: function of the junction’, Nature Reviews Molecular Cell Biology, 13, p. 607.

Ruggiero, A. et al. (2017) ‘Loss of forebrain MTCH2 decreases mitochondria motility and calcium handling and impairs hippocampal-dependent cognitive functions’, Scientific Reports, 7, p. 44401.

Santel, A. and Fuller, M.T. (2001) ‘Control of mitochondrial morphology by a human mitofusin’, Journal of Cell Science, 114(5), pp. 867 LP–874.

Schindelin, J. et al. (2012) ‘Fiji: an open-source platform for biological-image analysis’, Nature Methods, 9(7), pp. 676–682. doi:10.1038/nmeth.2019.

Sesaki, H. and Jensen, R.E. (1999) ‘Division versus fusion: Dnm1p and Fzo1p antagonistically regulate mitochondrial shape’, The Journal of cell biology, 147(4), pp. 699–706. doi:10.1083/jcb.147.4.699.

Tondera, D. et al. (2009) ‘SLP-2 is required for stress-induced mitochondrial hyperfusion.’, The EMBO journal, 28(11), pp. 1589–1600. doi:10.1038/emboj.2009.89.

Wydysh, E.A. et al. (2009) ‘Design and Synthesis of Small Molecule Glycerol 3-Phosphate Acyltransferase Inhibitors’, Journal of Medicinal Chemistry, 52(10), pp. 3317–3327. doi:10.1021/jm900251a.

Yamashita, A. et al. (2014) ‘Glycerophosphate/Acylglycerophosphate acyltransferases.’, Biology, 3(4), pp. 801–830. doi:10.3390/biology3040801.

Zaltsman, Y. et al. (2010) ‘MTCH2/MIMP is a major facilitator of tBID recruitment to mitochondria.’, Nature cell biology, 12(6), pp. 553–62. doi:10.1038/ncb2057.

Zhang, Y. et al. (2016) ‘Mitoguardin Regulates Mitochondrial Fusion through MitoPLD and Is Required for Neuronal Homeostasis’, Molecular Cell, 61(1), pp. 111–124. doi:https://doi.org/10.1016/j.molcel.2015.11.017.

Zhao, J. et al. (2011) ‘Human MIEF1 recruits Drp1 to mitochondrial outer membranes and promotes mitochondrial fusion rather than fission’, The EMBO journal, 30(14), pp. 2762–2778. doi:10.1038/emboj.2011.198.

